# Predicting binding free energies: Frontiers and benchmarks

**DOI:** 10.1101/074625

**Authors:** David L. Mobley, Michael K. Gilson

## Abstract

Binding free energy calculations based on molecular simulations provide predicted affinities for biomolecular complexes. These calculations begin with a detailed description of a system, including its chemical composition and the interactions between its components. Simulations of the system are then used to compute thermodynamic information, such as binding affinities. Because of their promise for guiding molecular design, these calculations have recently begun to see widespread applications in early stage drug discovery. However, many challenges remain to make them a robust and reliable tool. Here, we briefly explain how the calculations work, highlight key challenges, and argue for the development of accepted benchmark test systems that will help the research community generate and evaluate progress.

**Manuscript version 1.1.1 pre-release** See https://github.com/mobleylab/benchmarksets for all versions.

## I. INTRODUCTION

Molecular simulations provide a powerful technique for predicting and understanding the structure, function, dynamics, and interactions of biomolecules. Often, these techniques are valued because they provide a movie of what might be going on at the atomic level. However, simulations also can be used to make quantitative predictions of thermodynamic and kinetic properties, with applications in fields including drug discovery, chemical engineering, and nanoengineering. A thermodynamic property of particular interest is the binding affinity between biomolecules and ligands such as inhibitors, modulators, or activators. With accurate and rapid affinity predictions, we could use simulations in varied health-related applications, such as the prediction of biomolecular interaction networks in support of systems biology, or rapid design of new medications with reduced side-effects and drug resistance.

### A. Imagining a tool for drug discovery

A major aim in the development of molecular simulations is to create quantitative, accurate tools which will guide early stage drug discovery. Consider a medicinal chemist in the not-too-distant future who has just finished synthesizing several new derivatives of an existing inhibitor as potential drug leads targeting a particular biomolecule, and has obtained binding affinity or potency data against the desired biomolecular target. Before leaving work, he or she generates ideas for perhaps 100 new compounds which could be synthesized next, then sets a computer to work overnight prioritizing them. By morning, the compounds have all been prioritized based on reliable predictions of their affinity for the desired target, selectivity against alternative targets which should be avoided, solubility, and membrane permeability. The chemist then looks through the predicted properties for the top few compounds and selects the next ones for synthesis. If synthesizing and testing each compound takes several days, this workflow compresses roughly a year's work into a few days.

While this workflow is not yet a reality, huge strides have been made in this direction, with calculated binding affinity predictions now showing real promise [25, 26, 34, 38, 121, 155, 180, 187], solubility predictions beginning to come online [103, 138, 152], and predicted drug resistance/selectivity also apparently tractable [100], with some headway apparent on membrane permeability [30, 96]. A considerable amount of science and engineering still remains to make this vision a reality, but, given recent progress, the question now seems more one of *when* rather than *whether*.

### B. Increasing accuracy will yield increasing payoffs

Recent progress in computational power, especially the widespread availability of graphics processing units (GPUs) and advances in automation [105] and sampling protocols, have helped simulation-based techniques reach the point where they now appear to have sufficient accuracy to be genuinely useful in guiding pharmaceutical drug discovery, at least for a certain subset of problems [25, 34, 77, 117, 155, 180, 187]. Specifically, in some situations, free energy calculations appear to be capable of achieving RMS errors in the 1-2 kcal/mol range with current force fields, even in prospective applications. As a consequence, pharmaceutical companies are beginning to use these methods in discovery projects. The most immediate application of these techniques is to guide synthesis for lead optimization, but applications to scaffold hopping and in other areas also appear possible.

At the same time, it is clear that not all situations are so favorable, so it is worth asking what level of accuracy is actually needed. It is often suggested that we need binding free energy predictions accurate to within ∼ 1 kcal/mol, but we are not aware of a clear basis for this figure beyond the fact it is a pleasingly round number that is close to the thermal kinetic energy, *RT*. Instead of setting a single threshold requirement for accuracy, it is more informative to consider how accurate calculations must be to reduce the number of compounds synthesized and tested by some factor, relative to the number required without computational prioritization. If one targets a three-fold reduction, the answer appears to be that calculations with a 2 kcal/mol RMS error will suffice [121, 161]. Thus, one can gain substantial benefit from simulations that are good yet still quite imperfect.

More broadly, though, this analysis does not address the net value of computational affinity predictions in drug discovery. Costs include those of the software, computer time, and personnel required to incorporate calculations into the workflow; while benefits include the savings, revenue gains, and externalities attributable to reducing the number of low-affinity compounds synthesized and arriving earlier at a potent drug candidate. In addition, with sufficiently reliable predictions, chemists may choose to tackle difficult synthesis efforts they otherwise might have avoided, resulting in more novel and valuable chemical matter.

### C. Overview of free energy calculations

The present review focuses on a class of methods in which free energy differences are computed with simulations that sample Boltzmann distributions of molecular configurations. These samples are usually generated by molecular dynamics (MD) simulations [88], with the system effectively coupled to a heat bath at constant temperature, but Monte Carlo methods may also be used [29, 113, 114]. In either case, the energy of a given configuration is provided by a potential function, or force field, which estimates the potential energy of a system of solute and solvent molecules as a function of the coordinates of all of its atoms. Such simulations may be used in several different ways to compute binding free energies or relative binding free energies, as detailed elsewhere [24, 27, 114, 160] and summarized below. In all cases, however, the calculations yield the free energy difference between two states of a molecular system, and they do so by computing the reversible work for changing the initial state to the final one. Two broad approaches deserve mention.

The first general approach directly computes the standard free energy of binding of two molecules via evaluating the reversible work of transferring the ligand from the binding site into solution. (This is sometimes called an absolute binding free energy calculation.) The pathway of this change may be one that is physically realizable, or one that is only realizable *in silico*, in which case it is sometimes called an “alchemical” pathway. Physical pathway methods provide the standard binding free energy by computing the reversible work of, in effect, pulling the ligand out of the binding site. Although, by definition, the pathway used must be a physical one that could occur in nature, it need not be probable, and improbable pathways, governed by an order parameter specifying how far the ligand is from the binding site, are often used [11, 73, 78, 179, 192, 200]. In addition, artificial restraints may be useful to avoid sampling problems in the face of often complex barriers along the pathway [11, 73, 78, 179, 192]. By contrast, alchemical pathway methods artificially decouple the ligand from the binding site and then recouple it to solution from the protein [12, 66, 74, 87, 118]. Although alchemical decoupling methods may avoid clashes of the ligand with the protein that might be problematic when pathway methods are applied to a protein with a buried binding site, they still can pose some of the same sampling challenges. For example, sampling of the unbound receptor must be adequate after the ligand is removed, and water molecules must have time to equilibrate in the vacated binding site. Given that free energy is a state function, it is not surprising that alchemical and physical pathway approaches yield apparently comparable results when applied to the same systems [37, 71, 99, 199].

The second general approach computes the difference between the binding free energies of two different ligands for the same receptor, by computing the work of artificially converting one ligand into another, first in the bound state and then free in solution [24, 27, 114, 175]. Because these conversions are not physically realizable, such calculations are, again, called alchemical. These calculations can be quite efficient if the two ligands are very similar to each other, but they become more complicated and pose greater sampling problems if the two ligands are very different chemically or if there is a high barrier to interconversion between their most stable bound conformations [105]. In addition, there may be concerns about slow conformational relaxation of the protein in response to the change in ligand. Nonetheless, alchemical relative free energy calculations currently are the best automated and most widely used free energy methods [105, 121, 187].

The accuracy and precision of all of these methods are controlled by three major considerations. First, many conformations typically need to be generated, or sampled, in order to obtain an adequate representation of the Boltzmann distribution. In the limit of infinite sampling, a correctly implemented method would yield the single value of the free energy difference dictated by the specification of the molecular system and the chosen force field. In reality, however, only finite sampling is possible, so the reported free energy will differ from the nominal value associated with infinite sampling. In addition, because sampling methods are typically stochastic and the dynamics of molecular systems are sensitive to initial conditions [3], repeated calculations, using different random number seeds or initial states, will yield somewhat different results. The problem of finite sampling is most acute for systems where low-energy (hence highly occupied) conformational states are separated by high effective barriers, whether energetic or entropic. Second, even if adequate sampling is achievable, free energy differences may disagree substantially with experiment if the force field is not sufficiently accurate. Third, errors may also arise if the representation of the system in the simulation does not adequately represent the actual system, e.g. if protonation states are assigned incorrectly and held fixed.

### D. Challenges and the domain of applicability

Thus, in order for a free energy calculation to be reliable, it must use an appropriate representation of the physical system and an accurate force field, and it must adequately sample the relevant molecular configurations. In the case of the more widely used alchemical relative free energy approach, this means that the best results are expected when:

- a high quality receptor structure is available, without missing loops or other major uncertainties
- the protonation state of the ligand and binding-site residues (as well as any other relevant residues) can reliably be inferred
- the ligand binding mode is defined by crystallographic studies and is not expected to change much on modification
- the receptor does not undergo substantial, slow conformational changes
- key interactions are expected to be well-described by underlying force fields

Beyond this domain of applicability—whose dimensions are, in fact, still somewhat vague [1] — substantial challenges may be encountered. For example, binding free energy calculations for a cytochrome C peroxidase mutant suggest limitations of fixed-charge force fields. In this case, the strength of electrostatic interactions in a buried, relatively nonpolar binding site appears to be overestimated by a conventional fixed-charge force field, likely due to underestimation of polarization effects [147]. Sampling problems are also common, with slow sidechain rearrangements and ligand binding mode rearrangements in model binding sites in T4 lysozyme posing timescale problems unless enhanced or biased sampling methods are carefully applied [14, 57, 82, 119, 120, 186]; and larger-scale protein motions induced by some ligands can also pose challenges [14, 101].

Although such problems need not prevent free energy calculations from being used, they can require specific adjustment of procedures and parameters based on experience and knowledge of the system at hand. Thus, a key challenge for the field is how to use insights from well-studied cases to enable automation and reduce the detailed knowledge of each system required to carry out high quality simulations.

Troubleshooting is also a major challenge. In most cases where calculations diverge substantially from experiment, the reason for the discrepancy is not apparent. Is the force field inaccurate? Would the results improve with more sampling? Were protonation states misassigned—or do they perhaps even change on binding? There might even be a software bug [43] or a human error in the use of the software. As a consequence, it is not clear what steps are most urgently needed to advance the field as a whole.

## II. THE NEED FOR WELL-CHOSEN BENCHMARK SYSTEMS

Although tests of individual free energy methods are not uncommon today [25, 34, 117, 180, 187], the use of nonoverlapping molecular systems and computational protocols makes it difficult to compare methods on a rigorous basis. In addition, few studies are designed to identify key sources of error and thereby focus future research and development. A few molecular systems have now emerged as *de facto* standards for general study (Section III). These selections result in part from two series of blinded prediction challenges (SAMPL [130], and CSAR [42] followed by D3R [60]), which have helped focus the computational chemistry community on a succession of test cases and highlighted the need for methodological improvements. However, broader adoption of a larger and more persistent set of test cases is needed. By coalescing around a compact set of benchmarks, well chosen to challenge and probe free energy calculations, practitioners and developers will be able to better assess and drive progress in binding free energy calculations.

### A. Benchmark types and applications

We envision two classes of benchmark cases: “hard” benchmarks, which are simple enough that well-converged results can readily be computed; and “soft” benchmarks, for which convincingly converged results cannot readily be generated, but which are still simple enough that concerted study by the community can delineate key issues that might not arise in the simpler “hard” cases. The following subsections provide examples of how hard and soft benchmark systems may be used to address important issues in free energy simulations.

#### 1. Hard benchmarks

a. *Systems to test software implementations and usage* It is crucial yet nontrivial to validate that a simulation package correctly implements and applies the desired methods [159], and benchmark cases can help with this. First, all software packages could be tested for their ability to generate correct potential energies for a single configuration of a specified molecular system and force field. These results should be correct to within rounding error and the precision of the physical constants used in the calculations [159]. Similarly, different methods and software packages should give consistent binding free energies when identical force fields are applied with identical simulation setups and compositions. The benchmark systems for such testing can be simple and easy to converge, and high precision free energies (e.g., uncertainty ≈ 0.1 kcal/mol) should serve as a reference. Test calculations should typically agree with reference results to within 95% confidence intervals, from established methods [51, 158], For this purpose, the correctly computed values need not agree with experiment; indeed, experimental results are unnecessary.
b. *Systems to check sampling completeness and efficiency* As discussed above, free energy calculations require thorough sampling of molecular configurations from the Boltzmann distribution dictated by the force field that is employed. This sampling is typically done by running molecular dynamics simulations, and for systems as large and complex as proteins, it is difficult to carry out long enough simulations. Calculations with inadequate sampling yield results that are imprecise, in the sense that multiple independent calculations with slightly different initial conditions will yield significantly different results, and these ill-converged results will in general be poor estimates of the ideal result obtained in the limit of infinite sampling. Advanced simulation methods have been developed to speed convergence [160, 172], but it is not always clear how various methods compare to one another. To effectively compare such enhanced sampling methods, we need benchmark molecular systems, parameterized with a force field that many software packages can use, that embody various sampling challenges, such as high dimensionality, and energetic and entropic barriers between highly occupied states, but which are just tractable enough that reliable results are available via suitable reference calculations. Again, experimental data are not required, as the main purpose is to test and compare the power of various enhanced sampling technologies.
c. *Systems to assess force field accuracy* Some molecular systems are small and simple enough that current technology allows thorough conformational sampling, and hence well converged calculations of experimental observables. This has long been feasible for liquids [86]; for example, it is easy to precisely compute the heat of vaporization of liquid acetone with one of the standard force fields. More recently, advances in hardware and software have made it possible to compute binding thermodynamics to high precision for simple molecular recognition systems [73], as further discussed below. In such cases, absent complications like uncertain protonation states, the level of agreement with experiment reports directly on the accuracy of the force field. Thus, simple molecular recognition systems with reliable experimental binding data represent another valuable class of benchmarks. Here, of course, experimental data are needed. Ideally, the physical materials will be fairly easy to obtain so that measurements can be replicated, and new experimental conditions, such as different temperatures and solvent compositions, can be explored.

#### 2. Soft benchmarks

a. *Systems to challenge conformational sampling techniques* Enhanced sampling techniques (Section II A 1 b), designed to speed convergence of free energy simulations, may not be adequately tested by any hard benchmark, because such systems are necessarily rather simple. Thus, despite the fact that reliable reference results are not available for soft benchmarks, they are still important for method comparisons. For example, it may become clear that some methods are better at sampling in systems with high energy barriers, and others in highdimensional systems with rugged energy surfaces. Developers should test methods on a standard set of benchmark systems for informative comparisons.
b. *Direct tests of protein-ligand binding calculations* Although it is still very difficult to convincingly verify convergence of many protein-ligand binding calculations, it is still important to compare the performance of various methods for these real-world challenges. Appropriate soft benchmarks are likely to be cases which are still not overly challenging, involving small proteins and simple binding sites. We propose defining a series of benchmark protein-ligand systems that systematically introduce specific challenges. In particular, they should exemplify none, one, two, or N of the following challenges, in various combinations:

1. Sampling challenges

a. Sidechains in the binding site rearrange on binding different ligands
b. Modest backbone conformational changes, such as loop motion
c. Large scale conformational changes, such as domain motions and allostery
d. Ligand binding modes change unpredictably with small chemical modifications
e. High occupancy water sites rearrange depending on bound ligand
2. System challenges

a. Protonation state of ligand and/or protein changes on binding
b. Multiple protonation states of the ligand and/or receptor are relevant, due to pKas near the experimental pH, or the presence of multiple relevant tautomers
c. Results are sensitive to buffer, salts or other environmental factors
3. Force field challenges

a. Strong electric fields suggest that omission of explicit electronic polarizability will limit accuracy
b. Ligands interact directly with metal ions
c. Ligands or co-factors challenge existing force fields
c. *Progression of soft benchmarks* We envision these more complex benchmark systems proceeding through stages, initially serving effectively as a playground where major challenges and issues are explored, documented, and become well-known. Eventually, some will become sufficiently well characterized and sampled that they become hard benchmarks.

### B. Applications and limitations of benchmark systems

Benchmark systems along the lines sketched above will allow potential advances in computational methods to be tested in a straightforward, reproducible manner. For example, force fields may be assessed by swapping new parameters, or even a new functional form, into an existing workflow to see the impact on accuracy for a hard benchmark test. Sampling methods may be assessed by using various enhanced sampling methods for either hard or soft sampling benchmarks, here without focusing on accuracy relative to experiment. And system preparation tools could be varied to see how different approaches to assigning protonation states, modeling missing loops, or setting initial ligand poses, affect agreement of receptorligand binding calculations with experiment—with the understanding that force field and sampling also play a role. Finally, comparisons across methods will be greatly facilitated by community acceptance of a set of standard cases: well-characterized and studied benchmarks utilized by the majority of developers and research groups, ideally on a routine basis.

At the same time, there is a possibility that that some methods will inadvertently end up tuned specifically to generate good results for the set of accepted benchmarks. In such cases, the results for systems outside the benchmark set might still be disappointing. This means the field will need to work together to develop a truly representative set of benchmarks. This potential problem can also be mitigated by sharing of methods to enable broader testing by non-developers, and by participation in blinded prediction challenges, such as SAMPL and D3R, which confront methods with entirely new challenge cases.

## III. CURRENT BENCHMARK SYSTEMS FOR BINDING PREDICTIONS

No molecular systems have yet been designated by the field as benchmarks for free energy calculations, but certain host molecules and designed binding sites in the enzyme T4 lysozyme have emerged as particularly helpful and widely studied test cases. Here, we describe these artificial receptors and propose specific host-guest and T4 lysozyme-ligand combinations as initial benchmark systems for free energy calculations. We also point to several additional hosts and small proteins that also have potential to generate useful benchmarks in the future (Section IV). We focus on cases where experimental data are available and add value, rather than cases chosen to test conformational sampling methods, for which experimental data are not required (Section II A).

### A. Host-guest benchmarks

Chemical hosts are small molecules, often comprising fewer than 100 non-hydrogen atoms, with a cavity or cleft that allows them to bind other compounds, called guests, with significant affinity. Hosts bind their guests via the same basic forces that proteins used to bind their ligands, so they can serve as simple test systems for computational models of noncovalent binding. Moreover, their small size, and, in many cases, their rigidity, can make it feasible to sample all relevant conformations, making for “hard” benchmarks as defined above (Section II A). Furthermore, experiments can often be run under conditions that make the protonation states of the host and guest unambiguous. Under these conditions, the level of agreement of correctly executed calculations with experiment effectively reports on the validity of the force field (Section II A 1 c). For a number of host-guest systems, the use of isothermal titration calorimetry (ITC) to characterize binding provides both binding free energies and binding enthalpies. Binding enthalpies can often also be computed to good numerical precision [73], so they provide an additional check of the validity of simulations. A variety of curated host-guest binding data is available on BindingDB at http://bindingdb.org/bind/HostGuest.jsp.

Hosts fall into chemical families, such that all members of each family share a major chemical motif, but individuals vary in terms of localized chemical substitutions and, in some families, the number of characteristic monomers they comprise. For example, all members of the cyclodextrin family are chiral rings of glucose monomers; family members then differ in the number of monomers and in the presence or absence of various chemical substituents. For tests of computational methods ultimately aimed at predicting protein-ligand binding affinities in aqueous solution, water soluble hosts are, arguably, most relevant. On the other hand, host-guest systems in organic solvents may usefully test how well force fields work in the nonaqueous environment within a lipid membrane. Here, we focus on two host families, the cucurbiturils [52, 124] and the octa-acids (more generally, Gibb deep cavity cavitands) [62, 76]. These have already been the subject of concerted attention from the simulation community, due in part to their use in the SAMPL blinded prediction challenges [130, 132, 199].

#### 1. Cucurbiturils

The cucurbiturils (Figure 1) are achiral rings of glycoluril monomers [52]. The first characterized family member, cucurbit[6]uril, has six glycoluril units, and subsequent synthetic efforts led to the five-, seven-, eight- and ten-monomer versions, cucurbit[n]uril (n=5,6,7,8,10) [104], which have been characterized to different extents. The n=6,7,8 variants accommodate guests of progressively larger size, but are consistent in preferring to bind guests with a hydrophobic core sized to fit snugly into the relatively nonpolar binding cavity, along with at least one cationic moiety (though neutral compounds do bind [97, 194]) that forms stabilizing interactions with the oxygens of the carbonyl groups fringing both portals of the host [104]. Although derivatives of these host molecules have been made [4, 31, 98, 182], most of the binding data published for this class of hosts pertain to the non-derivatized forms. A fairly extensive set of data is available in BindingDB at http://bindingdb.org/bind/HostGuest.jsp.

**Fig. 1.**
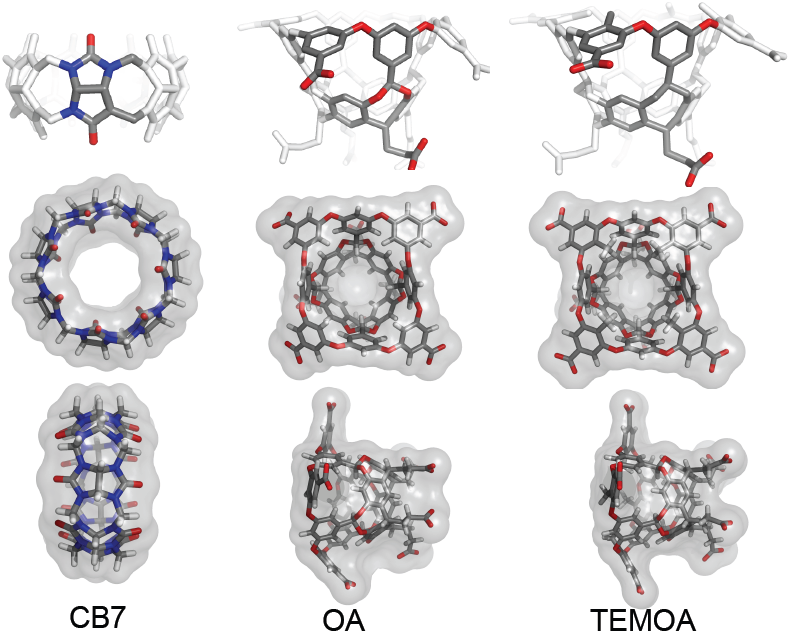
OA, TEMOA, and CB7 hosts. Shown are the hosts which are the focus of our host-guest benchmark sets – two variants of the octa-acid GDCC, and CB7, a cucurbituril. Guest structures are available in the supplemental material.

We propose cucurbit[7]uril (CB7) as the basis of one series of host-guest benchmark systems (Figure 1). This host is convenient experimentally, because it is reasonably soluble in water; and computationally, because it is quite rigid and lacks acidic or basic groups. In addition, it has attracted particular interest because of the high binding affinities of some guests, exceeding even the tightest-binding protein-ligand systems [19, 104, 125, 146]. Finally, CB7 is already familiar to a number of computational chemistry groups, as it figured in two of the three SAMPL challenges that included host-guest components [130, 132], and it is currently the focus of the “hydrophobe challenge” [153].

a. *CB7 presents several challenges* Despite the simplicity of CB7, calculations of its binding thermodynamics are still challenging, with several known complexities:

1. **Tight exit portal**: Guest molecules with bulky hydrophobic cores, such as adamantyl or [2.2.2]bicyclooctyl [125, 126] groups, do not fit easily through the constrictive portals [178]. As a consequence, free energy methods which compute the work of binding along a physical dissociation pathway may encounter a high barrier as the bulky core exits the cavity, and this can lead to subtle convergence problems [73, 179]. One way to solve this problem is to reversibly add restraints that open the portal, then remove the guest, and finally reversibly remove the restraints [73], including all of these contributions in the overall work of dissociation.
2. **Water binding and unbinding**: If one computes the work of removing the guest from the host by a nonphysical pathway, in which the bound guest is gradually decoupled from the host and surrounding water [66], large fluctuations in the number of water molecules within the host's cavity can occur when the guest is partly decoupled, and these fluctuations can slow convergence [150].
3. **Salt concentration and buffer conditions**: Binding thermodynamics are sensitive to the composition of dissolved salts, both experimentally [125, 126, 130] and computationally [78, 133]. As a consequence, to be valid, a comparison of calculation with experiment must adequately model the experimental salt conditions.
4. **Finite-size artifacts due to charge modification**: Because many guest molecules carry net charge, it should be ascertained that calculations in which guests are decoupled from the system do not generate artifacts related to the treatment of long-ranged Coulombic interactions [102, 144, 148, 163].
b. *The proposed CB7 benchmark sets comprise two compound series* For CB7, we have selected two sets of guests that were studied experimentally under uniform conditions (50 mM sodium acetate buffer, pH 4.74, 298K) by one research group [19, 104]. Each series is based on a common chemical scaffold, making it amenable to not only absolute but also alchemical relative free energy calculations (Section IC). One set is based on an adamantane core (Table I), and the other on an aromatic ring (Table II). These systems can be run to convergence to allow detailed comparisons among methods and with experiment. Their measured binding free energies range from −5.99 to −17.19 kcal/mol, with the adamantane series spanning a particularly large range.
c. *Prior studies provide additional insight into CB7's challenges* Sampling of the host appears relatively straightforward in CB7 as it is quite rigid and its symmetry provides for clever convergence checks [73, 127]. Due to its top-bottom symmetry, flips of guests from “head-in” to “head-out” configurations are not necessary to obtain convergence [47]. However, sampling of the guest geometry can be a challenge, with transitions between binding modes as slow as 0.07 flips/ns [127], and flexible guests also presenting challenges [127]. As noted above, water sampling can also be an issue, with wetting/dewetting transitions occurring on the 50 ns timescale [150] when the guest is partly decoupled from the aqueous host in alchemical calculations.

**Table I.**
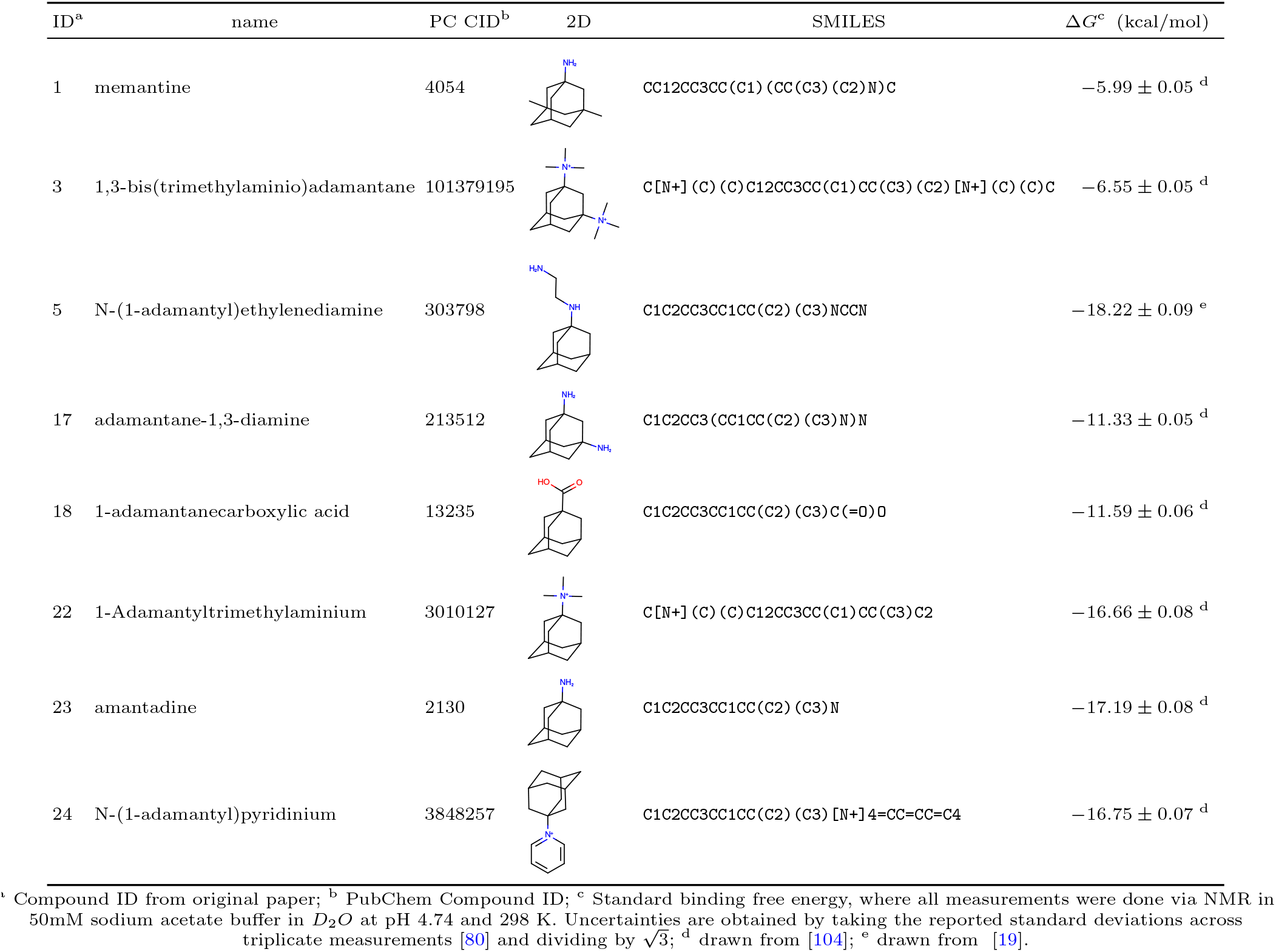
Proposed CB7 Set 1 benchmark data

| ID<sup>a</sup> | name | PC CID<sup>b</sup> | 2D | SMILES | $\Delta G^c$ (kcal/mol) |
| --- | --- | --- | --- | --- | --- |
| 1 | memantine | 4054 | | CC12CC3CC(C1)(CC(C3)(C2)N)C | $-5.99 \pm 0.05$ <sup>d</sup> |
| 3 | 1,3-bis(trimethylaminio)adamantane | 101379195 | | C[N+](C)(C)C12CC3CC(C1)CC(C3)(C2)[N+](C)(C)C | $-6.55 \pm 0.05$ <sup>d</sup> |
| 5 | N-(1-adamantyl)ethylenediamine | 303798 | | C1C2CC3CC1CC(C2)(C3)NCCN | $-18.22 \pm 0.09$ <sup>e</sup> |
| 17 | adamantane-1,3-diamine | 213512 | | C1C2CC3(CC1CC(C2)(C3)N)N | $-11.33 \pm 0.05$ <sup>d</sup> |
| 18 | 1-adamantanecarboxylic acid | 13235 | | C1C2CC3CC1CC(C2)(C3)C(=O)O | $-11.59 \pm 0.06$ <sup>d</sup> |
| 22 | 1-Adamantyltrimethylaminium | 3010127 | | C[N+](C)(C)C12CC3CC(C1)CC(C3)C2 | $-16.66 \pm 0.08$ <sup>d</sup> |
| 23 | amantadine | 2130 | | C1C2CC3CC1CC(C2)(C3)N | $-17.19 \pm 0.08$ <sup>d</sup> |
| 24 | N-(1-adamantyl)pyridinium | 3848257 | | C1C2CC3CC1CC(C2)(C3)[N+]4=CC=CC=C4 | $-16.75 \pm 0.07$ <sup>d</sup> |

<sup>a</sup> Compound ID from original paper; <sup>b</sup> PubChem Compound ID; <sup>c</sup> Standard binding free energy, where all measurements were done via NMR in 50mM sodium acetate buffer in $D_2O$ at pH 4.74 and 298 K. Uncertainties are obtained by taking the reported standard deviations across triplicate measurements [80] and dividing by $\sqrt{3}$ ; <sup>d</sup> drawn from [104]; <sup>e</sup> drawn from [19].

**Table II.**
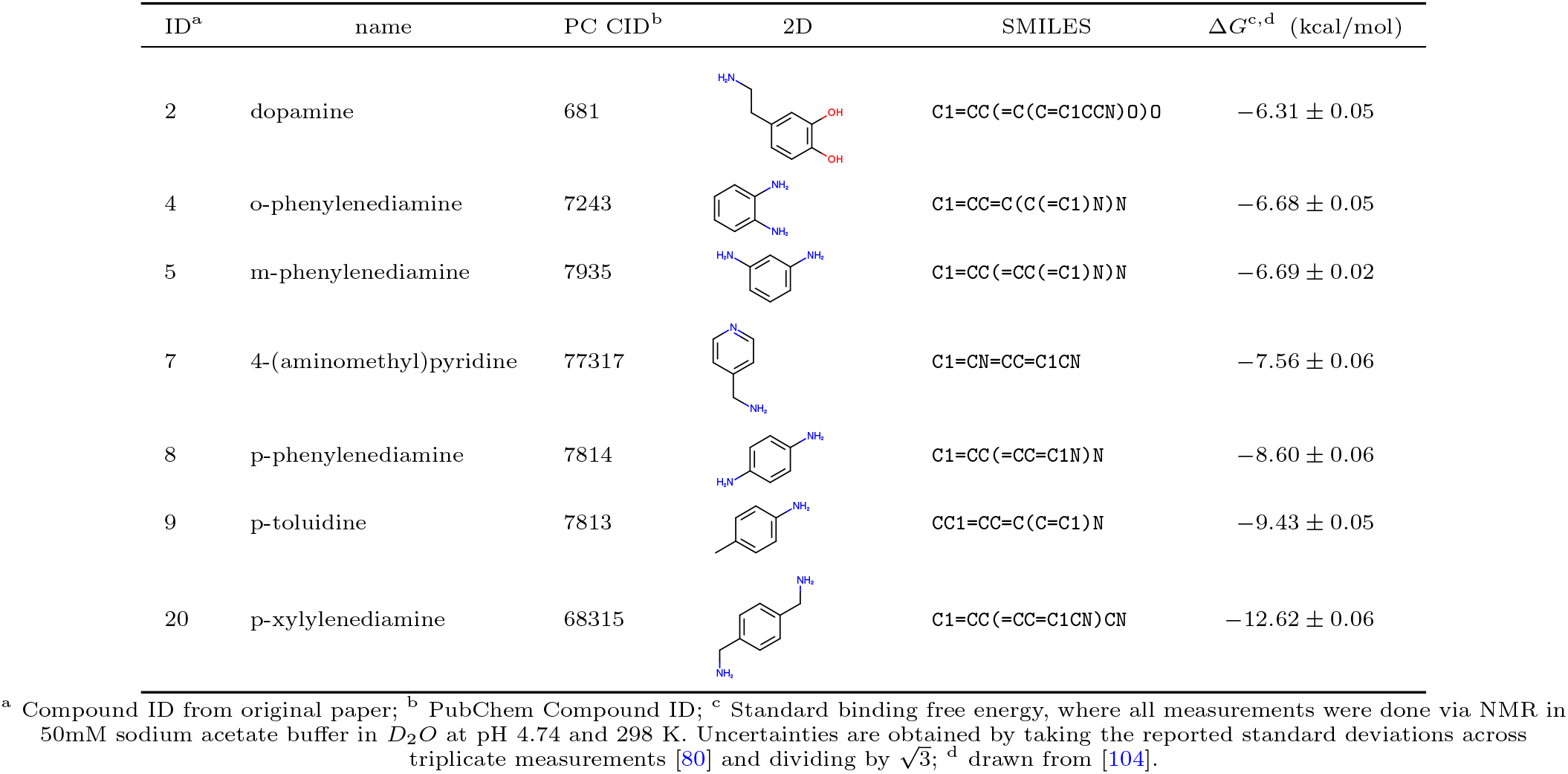
Proposed CB7 Set 2 benchmark data

Salt and buffer conditions are also key. In addition to the strong salt-dependence of binding [126], acetic acid (such as in a sodium acetate buffer) can compete with guests for the binding site [125]. This may partially explain systematic errors in some computational studies [78, 133]. Indeed, the difference between 50 mM sodium acetate buffer and 100 mM sodium phosphate buffer impacts measured binding free energies by 2.5-2.8 kcal/mol [130, 133]. Cationic guests could also have substantial and differing interactions with the counterions in solution, potentially lowering affinity relative to zero-salt conditions [130]. Additionally, CB7 can also bind cations fairly strongly [18, 79, 109]. Thus, one group found a 6.4-6.8 kcal/mol dependence on salt concentration [78] (possibly due to cation competition for the binding site), possibly impacting other studies as well [127].

Despite these issues, CB7 appears to be at the point where careful studies can probe the true accuracy of our force fields [59, 73, 78, 131, 197], and the results can be sobering, with RMS errors in the binding free energies as high as 8 kcal/mol [73, 127]. More encouragingly, the values of *R*^2^ values can be as high as 0.92 [73]. Calculated values are in many cases quite sensitive to details of force field parameters [126, 127, 131]. For example, modest modification of some Lennard-Jones parameters yielded dramatic improvements in calculated values [197], and host-guest binding data has, accordingly, been suggested as an input for force field development [59, 73, 197]. Water structure around CB7 and calculated binding enthalpies also appear particularly sensitive to the choice of water model [47, 59, 150], and water is clearly important for modulating binding [136]. The water model also impacts the number of sodium ions which must be displaced (in sodium-based buffer) on binding [59, 73].

In summary, CB7 is still a challenging benchmark that can put important issues into high relief. For example, in SAMPL4, free energy methods yielded *R*^2^ values from 0.1 to 0.8 and RMS errors of about 1.9 to 4.9 kcal/mol for the same set of CB7 cases [130]. This spread of results across rather similar methods highlights the need for shared benchmarks. Potential explanations include convergence difficulties, subtle methodological differences, and details of how the methods were applied. Until the origin of such discrepancies is clear, it is difficult to know how accurate our methods truly are.

#### 2. Gibb Deep Cavity Cavitands (GDCC)

The octa-acids (OA) (Figure 1) are synthetic hosts with deep, basket-shaped, hydrophobic binding sites [62]. The eight carboxylic acidic groups for which they were originally named make these hosts water-soluble, but do not interact directly with bound hosts; instead, they project outward into solvent. Binding data have been reported for the original form of this host (OA) [62] and for a derivative with four added methyl groups at equivalent locations in the entryway, where they can contact a bound guest (TEMOA) [58, 169]. (Note that OA and TEMOA have also been called OAH and OAMe, respectively [199].) Additional family members with other substituents around the portal have been reported, as has a new series in which the eponymic carboxylic groups are replaced by various other groups, including a number of basic amines [76]. However, we are not aware of binding data for these derivatives. Because these closely related hosts are clearly in the same family but do not have eight acidic groups, and in recognition of the family's developer, we propose the more general name Gibb deep cavity cavitands (GDCCs) for this family of hosts. The binding cavities of the GDCCs are fairly rigid, though less so than the cucurbiturils. Some simulators report “breathing” motions that vary the diameter of the entry by up to 8 Å[116]; and, in some studies, the benzoic acid “flaps” around the entry occasionally flip upward and into contact with the guest [176, 198], though this motion has not been verified experimentally. Additionally, the four priopionate groups protruding into solution from the exterior base of the cavity are all flexible.

The octa-acids tend to bind guest molecules possesisng a hydrophobic moiety that fits into the host's cavity and a hydrophilic moiety that projects into the aqueous solvent. Within these specifications, they bind a diversity of ligands, including both organic cations and anions, as well as neutral compounds with varying degrees of polarity [63, 65]. Compounds with adamantane or noradamantane groups display perhaps the highest affinities observed so far, with binding free energies ranging to about −8 kcal/mol [170]. Much of the experimental binding data comes from ITC, so binding enthalpies are often available.

Two experimental aspects of binding are particularly intriguing and noteworthy. First, the binding thermodynamics of OA is sensitive to the type and concentration of anions in solution. Although NaCl produces relatively modest effects, 100 mM sodium perchlorate, chlorate and isothiocyanate can shift binding enthalpies by up to about 10 kcal/mol and free energies by around 2 kcal/mol [64]. These effects are due in part to binding of anions by the host; indeed, trichloroacetate is reported to bind OA with a free energy of −5.2 kcal/mol [167], and competition of other guests with bound anions leads to entropy-enthalpy tradeoffs. Second, elongated guests can generate ternary complexes, in which two OA hosts encapsulate one guest, especially if both ends of the guest are not very polar [63].

a. *The proposed GDCC benchmark sets are drawn from SAMPL* We propose the establishment of two GDCC benchmark sets, based on data which formed part of the SAMPL4 and SAMPL5 challenges. One set is based on experiments carried out in phosphate buffer at pH 11.5, and the other on experiments in tetraborate buffer at pH 9.2 The guests in the first set (Table III) are adamantane derivatives and cyclic (aromatic and saturated) carboxylic acids (Table III), which bind to hosts OA and TEMOA with free energies of −3.7 to −7.6 kcal/mol. The second set of guests (Table IV) comprises carboxylic acids based on phenyl and cyclohexane cores. Both sets offer aqueous binding data with free energies spanning about 4 kcal/mol, frequently along with binding enthalpies. The hosts and many or all of their guests are small and rigid enough to allow convincing convergence of binding thermodynamics with readily feasible simulations; and, like the cucurbiturils, they are already emerging as *de facto* computational benchmarks, due to their use in the SAMPL4 and SAMPL5 challenges [130, 199].
b. *The GDCC hosts introduce new challenges beyond CB7* Issues deserving attention when interpreting the experimental data and calculating the binding thermodynamics of these systems include the following:

1. **Tight exit portal**: The methyl groups of the TEMOA variant narrow the entryway and can generate a barrier to the entry or exit of guest molecules with bulky hydrophobic cores, though the degree of constriction is not as marked as for CB7 (above). The TEMOA methyl groups can additionally hinder sampling of guest poses in the bound state, leading to convergence problems [199]
2. **Host conformational sampling**: Although the flexible propionate groups are not proximal to the binding cavity, they are charged and so can have long-ranged interactions. As a consequence, it may be important to ensure their conformations are well sampled, though motions may be slow [116]. Similarly, benzoic acid flips [176, 198] could potentially be an important challenge in some force fields.
3. **Water binding and unbinding**: Water appears to undergo slow motions into and out of the OA host, on timescales upwards of 5 ns [45]. This poses significant challenges for some approaches, such as metadynamics, where deliberately restraining water to stay out of the cavity when the host is not bound (and computing the free energy of doing so) can help convergence [11], and perhaps for other methods as well.
4. **Salt concentration and buffer conditions**: As in the case of CB7, binding to GDCCs is modulated by the composition of dissolved salts, both experimentally [64, 167] and computationally [137, 176]. As a consequence, to be valid, a comparison of calculation with experiment must adequately model the experimental salt conditions.
5. **Finite-size artifacts due to charge modification**: As for CB7, it should be ascertained that calculations in which charged guests are decoupled from the system do not generate artifacts related to long range Coulomb interactions. [102, 144, 148, 163].
6. **Protonation state effects**: Although experiments are typically run at pH values that lead to well-defined protonation states of the host and its guests, this may not always hold [45, 130, 176], particularly given experimental evidence for extreme binding-driven pKa shifts of 3-4 log units for some carboxylate compounds [167, 185]. Thus, attention should be given to ionization states and their modulation by binding.
c. *Prior studies provide additional insight into the challenges of OA* As noted, two different host conformational sampling issues have been observed, with dihedral transitions for the proprionate groups occurring on 1-2 ns timescales [116]); motions of the benzoic acid flaps were also relatively slow [176, 198] though perhaps thermodynamically unimportant. Guest sampling can also be an issue, at least in TEMOA [199], and this host's tight cavity may also have implications for binding entropy [198].

**Table III.**
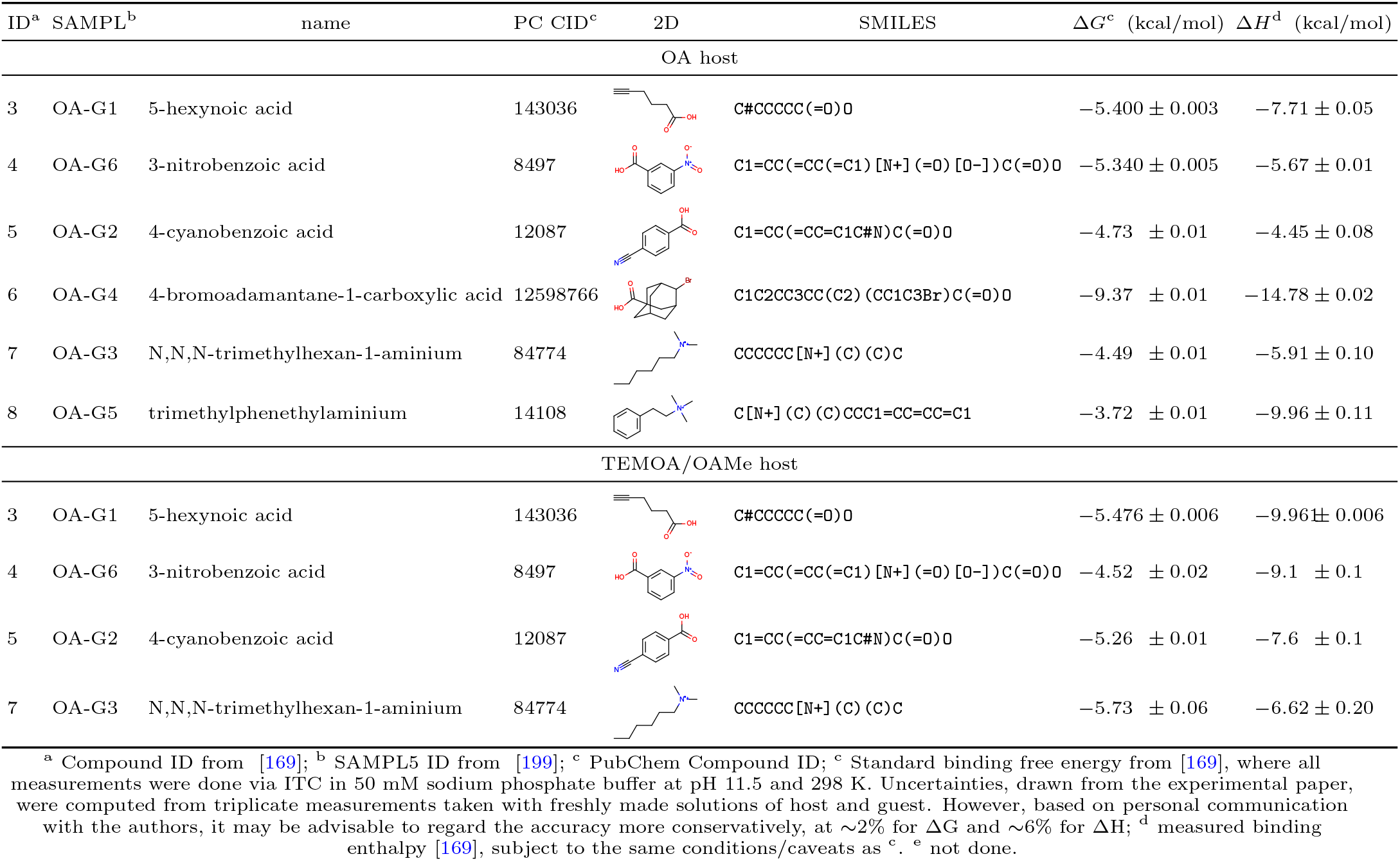
Proposed GDCC Set 1 benchmark data

**Table IV.**
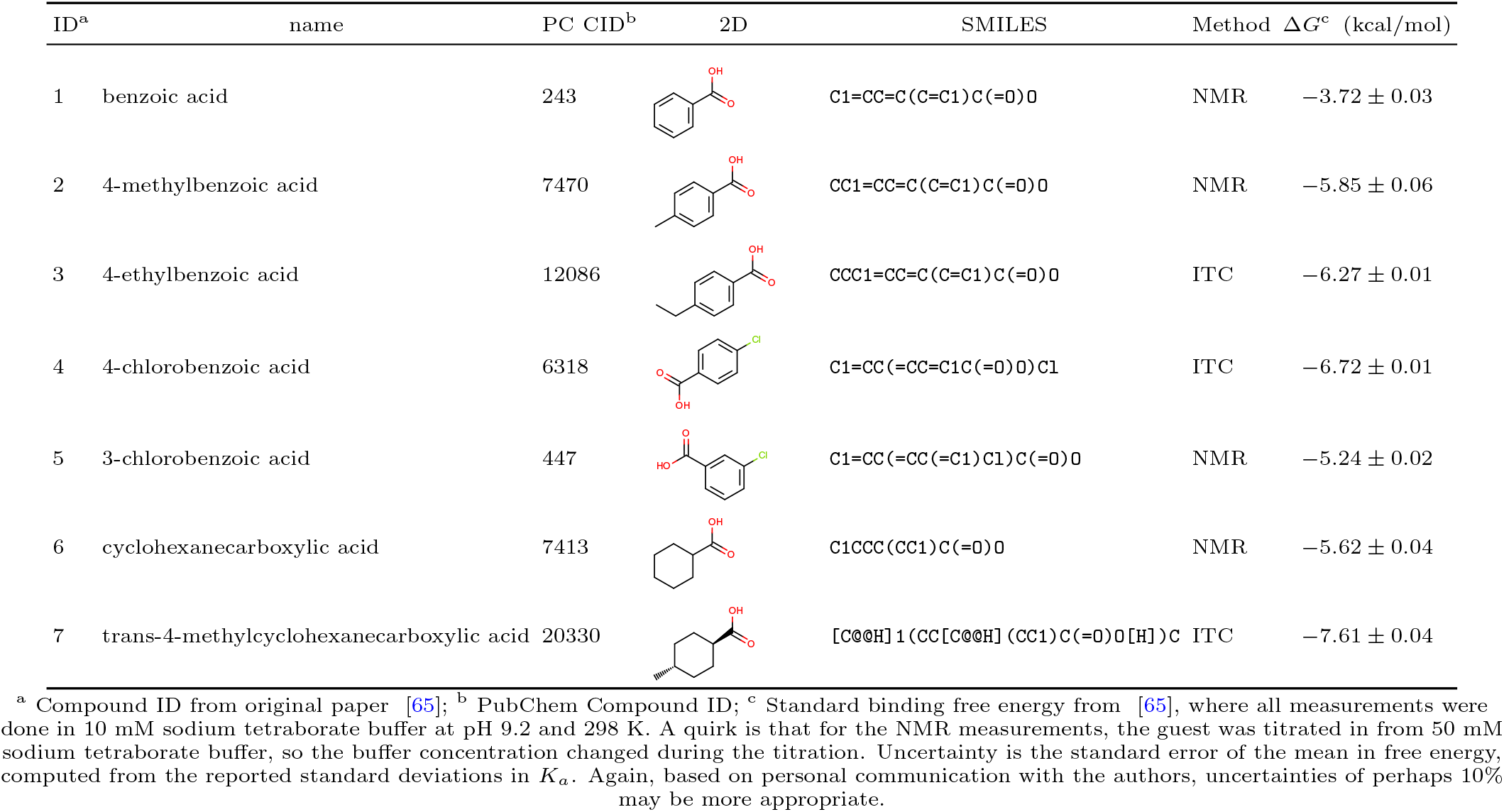
Proposed GDCC Set 2 benchmark data for binding to the OA host.

Salt concentration strongly modulates binding affinity, at least for anions, and the nature of the salt also plays an important role [20]. Co-solvent anions can also increase *or* decrease binding depending on their identity [64]. Some salts even bind to OA themselves, with perchlorate [64] and trichloroacetate [167] being particularly potent, and thus will compete with guests for binding. Computationally, including additional salt beyond that needed for system neutralization changed binding free energies by up to 4 kcal/mol [176].

Although the protonation states of the guests might seem clear and unambiguous, experimental evidence also indicates major pKa shifts on binding so that species such as acetate, formate and others could bind in neutral form at neutral pH [167, 185]. One study used absolute binding free energy calculations for different guest charge states, coupled with pKa calculations, and found that inclusion of pKa corrections and the possibility of alternate charge states of the guests affected calculated binding free energies by up to 2 kcal/mol [176]. Even the host protonation state may be unclear. Although OA might be assumed to have all eight carboxylic acids deprotonated at the basic pH of typical experiments, the four at the bottom are in close proximity, and these might make hydrogen bonds allowing retention of two protons [45]. Thus, there are uncertainties as to the host protonation state [45, 130], which perhaps also could be modulated by guest binding.

The challenges of system definition and conformational sampling appear to be greater for OA and TEMOA than for CB7, as discussed above, so it is harder to draw definite conclusions regarding force field accuracy and differences among force fields. Thus, although there have been some comparisons of charge [116, 127, 130] and water models [198], the resulting differences in computed binding thermodynamics so far seem mostly inconclusive or at least not that large. Similarly, OPLS and GAFF results did not appear dramatically different in accuracy [11]. It is worth noting that several groups using different computational approaches but the same force field and water model in SAMPL5 did not obtain identical binding free energies [11, 13, 199]. Some of these issues were resolved in follow-up work [11], bringing the methods into fairly good agreement for the majority of cases [13, 198].

### B. Protein-ligand benchmarks: the T4 lysozyme model binding sites

Although we seek ultimately to predict binding in systems of direct pharmaceutical relevance, simpler proteinligand systems can represent important stepping stones in this direction. Two model binding sites in T4 lysozyme have been particularly useful in this regard (Figure 2). These two binding sites, called L99A [128, 129] and L99A/M102Q [69, 188] for point mutations which create the cavities of interest, have been studied extensively experimentally and via modeling. As proteinligand systems, they introduce additional complexities beyond those observed in host-guest systems, yet they share some of the same simplicity. The ligands are generally small, neutral, and relatively rigid, with clear protonation states. In many cases, substantial protein motions do not occur on binding, helping calculated binding free energies to reach apparent convergence relatively easily. However, like host-guest systems, these binding sites are still surprisingly challenging [14, 57, 82, 101, 118–120]. Thus, precise convergence is sometimes difficult to achieve, and it is in all cases essentially impossible to fully verify. As a consequence, these are “soft benchmarks” as defined above (Section II A). The utility of the lysozyme model sites is also driven by the large body of available experimental data. It has been relatively easy to identify new ligands and obtain high quality crystal structures and affinity measurements, and this has allowed two different rounds of blinded free energy prediction exercises [14, 120].

**Fig. 2.**
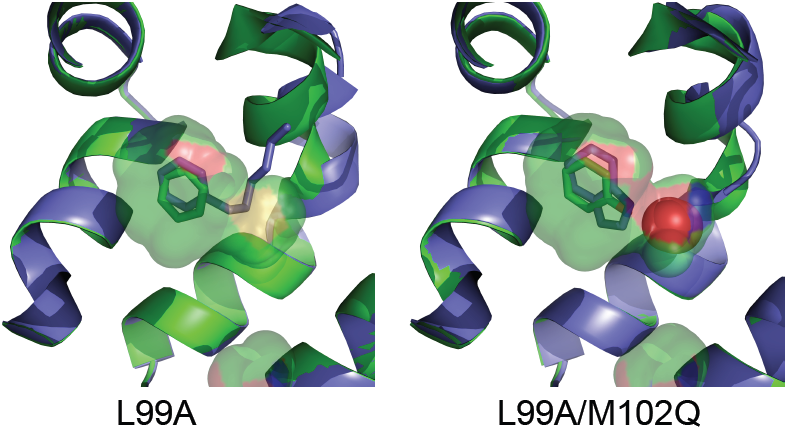
Benzene and hexylbenzene in the apolar lysozyme L99A site (left), and phenol and 4,5,6,7-tetrahydroindole in the polar L99A/M102Q site (right). These structures are from PDBs 4W52, 4W59, 1LI2, and 3HUA, respectively. The binding site cavities are shown as a semi-transparent surface, and the protein is shown with cartoons. In both cases, the structure with the smaller ligand is shown in green and that with the larger ligand is shown in blue; the larger ligand (hexylbenzene in L99A, 4,5,6,7-tetrahydroindole in L99A/M102Q) induces a motion of helix F bordering the binding site. Phenol and 4,5,6,7-tetrahydroindole both also bind with an ordered water (red sphere, right panel), though this does not occur for all ligands in the polar L99A/M102Q site.

#### 1. The apolar and polar cavities and their ligands

The L99A site is also called the “apolar” cavity. It is relatively flat and elongated, and binds mostly nonpolar molecules such as benzene, toluene, p-xylene, and n-butylbenzene: basically, a fairly broad range of nonpolar planar five- and six-membered rings and ring systems (such as indole). The polar version, L99A/M102Q, introduces an additional point mutation along one edge of the binding site, providing a glutamine that introduces polarity and the potential for hydrogen bonding. It still binds a variety of nonpolar ligands such as toluene (though not benzene). One small downside of these binding sites is that the range of affinities is relatively narrow: about −4.5 to −6.7 kcal/mol in the apolar site [120, 128], and about −4 to −5.5 kcal/mol in the polar site [14]. Thus, even the strongest binders are not particularly strong, and the weakest binders tend to run up against their solubility limits. Still, these sites offer immensely useful tests for free energy calculations.

For both sites, fixed charge force fields seem to yield reasonably accurate free energies, with RMS errors of 1-2 kcal/mol, and some level of correlation with experiment, despite limited dynamic range [14, 39, 57, 120, 184]. System composition/preparation issues also do not seem to be a huge factor. Instead, sampling issues predominate:

1. Ligand orientation: These oblong binding sites are buried, and their ligands are similar in shape. Ligands with axial symmetry typically have at least two reasonably likely binding modes, but broken symmetry can drive up the number of likely binding modes. For example, phenol has two plausible binding modes in the polar cavity [14, 70] but 3-chlorophenol has at least four, three of which appear to have some population in simulations [57], because the chlorine could point in either direction within the site. Timescales for binding mode interconversion are relatively slow, with in-plane transitions on the 1-10 nanosecond timescale, and out-of-plane transitions (e.g. between toluene's two symmetry-equivalent binding modes) taking hundreds of nanoseconds (Mobley group, unpublished data).
2. Sidechain rearrangement: Some sidechains reorganize when certain ligands, particularly larger ones, bind. This is common in particular for the side chain of Val 111 in the L99A site [82, 119, 129] and Leu 118, Val 111, and Val 103 in L99A/M102Q [14, 70, 188, 189]. These sidechain motions can be slow, due to the tight packing of the binding site, and therefore can present sampling problems for standard MD simulations [14, 82, 119, 120, 186].
3. Backbone rearrangement: Larger ligands induce shifts of the F helix (residues 107 or 108 to 115), which is adjacent to the binding site, allowing the site to enlarge. This occurs in both binding sites [14, 111, 189], but is best characterized for L99A [111]. There, addition of a series of methyl groups from benzene up to n-hexylbenzene causes a conformational transition in the protein from closed to intermediate to open conformations.

Tables V and VI introduce two initial benchmark sets based on the apolar and polar T4 lysozyme binding sites. These sets include ligands amenable to both absolute and relative free energy calculations, and have affinities that cover the currently available experimental range. Co-crystal structures are available in most cases; the corresponding PDB IDs are provided in the tables. The selected ligands span a range of challenges and levels of difficulty, ranging from fairly simple to including most of the challenges noted above. Essentially all of them have been included in at least one prior computational study, and some have appeared in a variety of prior studies. Additional known ligands and nonbinders are available, with binding affinities available for 19 compounds in the L99A site [44, 120, 128] and 16 in L99A/M102Q [14, 69, 188]. Because of the extent of the sampling challenges in lysozyme, binding of most ligands will currently constitute a soft benchmark, though long-timescale simulations to turn these into hard benchmarks may already be feasible.

**Table V.**
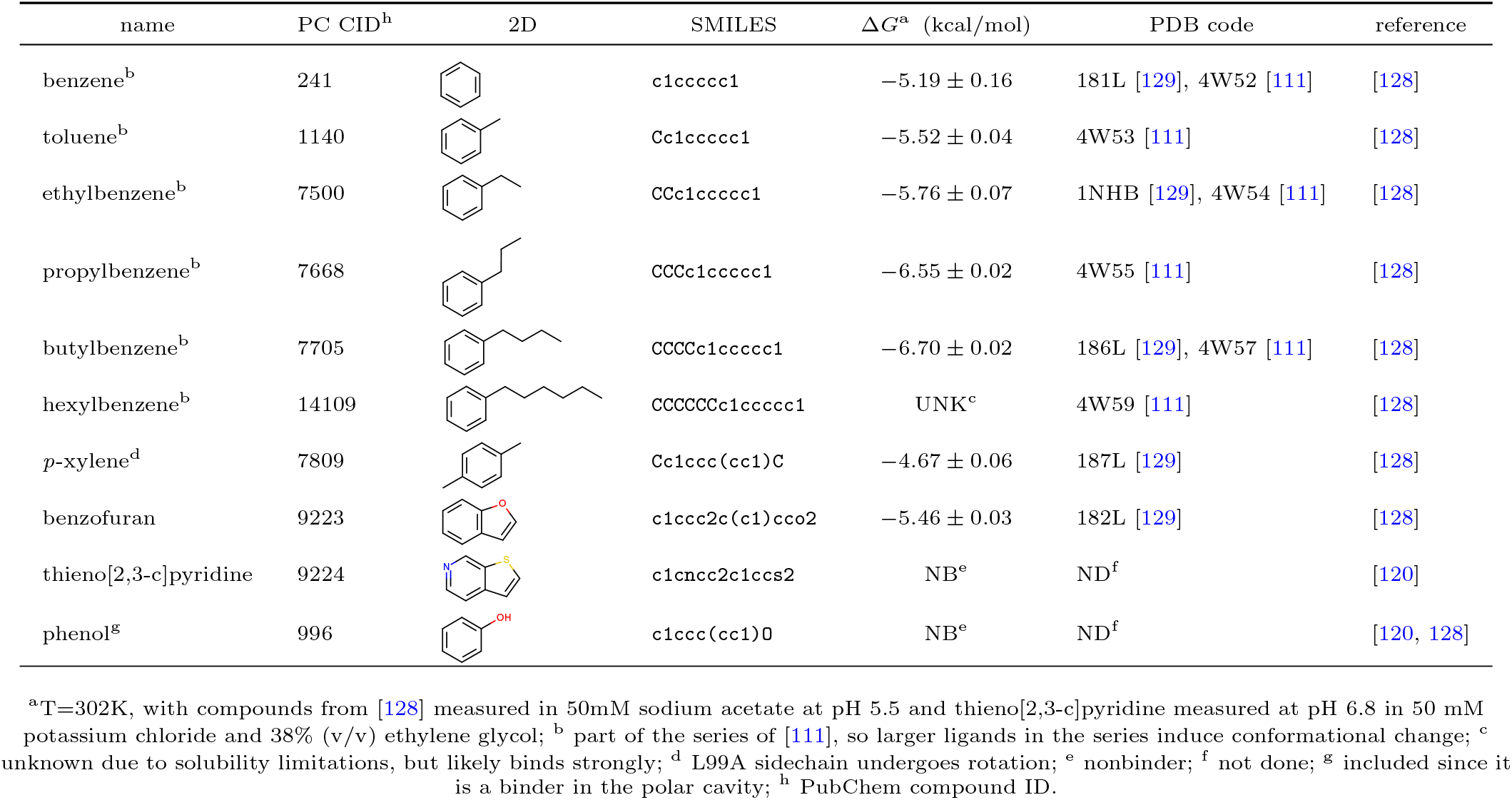
Proposed Lysozyme L99A Set benchmark data

| name | PC CID<sup>b</sup> | 2D | SMILES | $\Delta G^a$ (kcal/mol) | PDB code | reference |
| --- | --- | --- | --- | --- | --- | --- |
| benzene<sup>b</sup> | 241 | | c1ccccc1 | $-5.19 \pm 0.16$ | 181L [129], 4W52 [111] | [128] |
| toluene<sup>b</sup> | 1140 | | Cc1ccccc1 | $-5.52 \pm 0.04$ | 4W53 [111] | [128] |
| ethylbenzene<sup>b</sup> | 7500 | | CCc1ccccc1 | $-5.76 \pm 0.07$ | 1NHB [129], 4W54 [111] | [128] |
| propylbenzene<sup>b</sup> | 7668 | | CCCc1ccccc1 | $-6.55 \pm 0.02$ | 4W55 [111] | [128] |
| butylbenzene<sup>b</sup> | 7705 | | CCCCc1ccccc1 | $-6.70 \pm 0.02$ | 186L [129], 4W57 [111] | [128] |
| hexylbenzene<sup>b</sup> | 14109 |  | CCCCCCc1ccccc1 | UNK<sup>c</sup> | 4W59 [111] | [128] |
| *p*-xylene<sup>d</sup> | 7809 | | Cc1ccc(cc1)C | $-4.67 \pm 0.06$ | 187L [129] | [128] |
| benzofuran | 9223 | | c1ccc2c(c1)cco2 | $-5.46 \pm 0.03$ | 182L [129] | [128] |
| thieno[2,3-*c*]pyridine | 9224 |  | c1cncc2c1ccs2 | NB<sup>e</sup> | ND<sup>f</sup> | [120] |
| phenol<sup>g</sup> | 996 |  | c1ccc(cc1)O | NB<sup>e</sup> | ND<sup>f</sup> | [120, 128] |

<sup>a</sup>T=302K, with compounds from [128] measured in 50mM sodium acetate at pH 5.5 and thieno[2,3-*c*]pyridine measured at pH 6.8 in 50 mM potassium chloride and 38% (v/v) ethylene glycol; <sup>b</sup> part of the series of [111], so larger ligands in the series induce conformational change; <sup>c</sup> unknown due to solubility limitations, but likely binds strongly; <sup>d</sup> L99A sidechain undergoes rotation; <sup>e</sup> nonbinder; <sup>f</sup> not done; <sup>g</sup> included since it is a binder in the polar cavity; <sup>h</sup> PubChem compound ID.

**Table VI.**
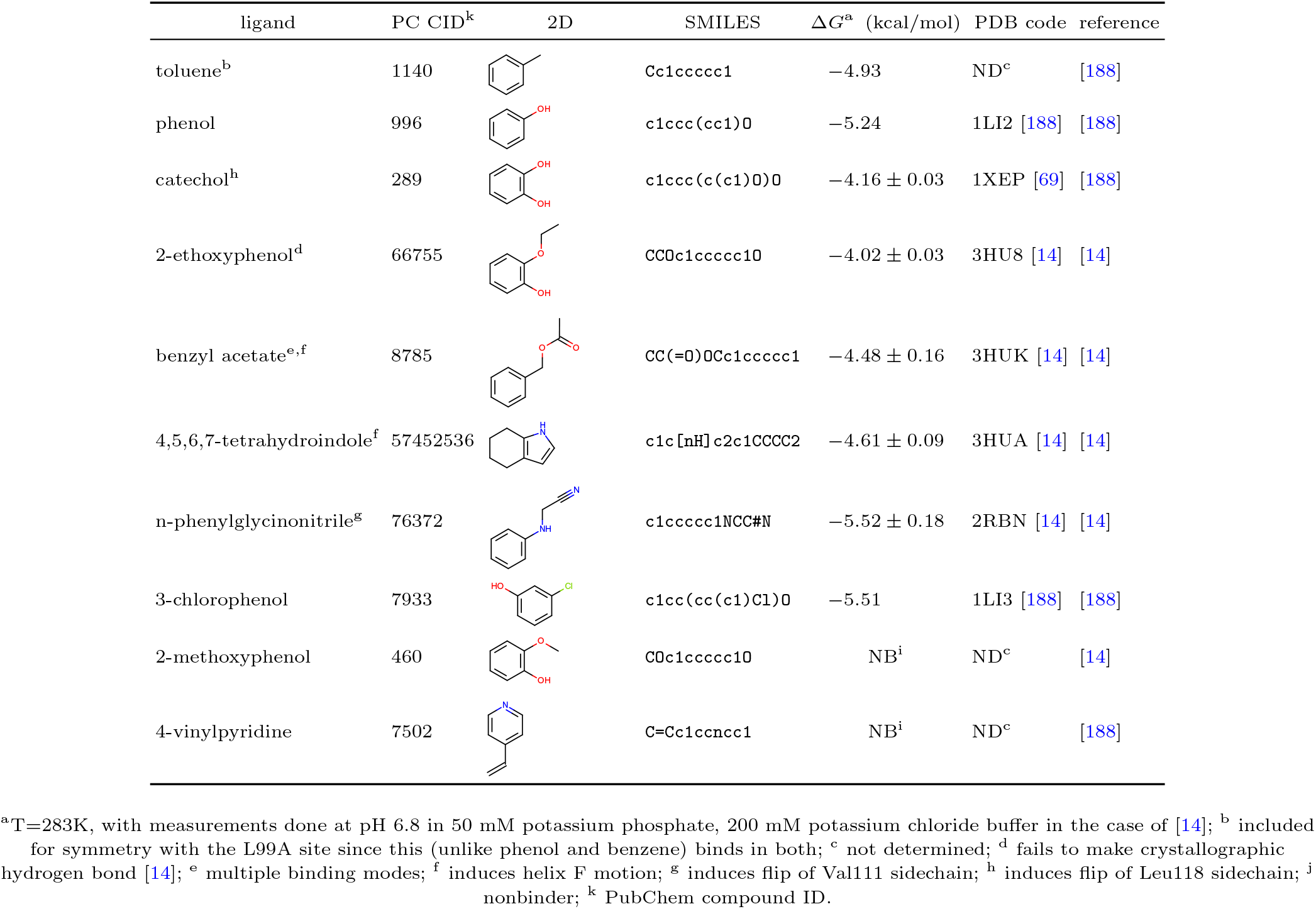
Proposed Lysozyme L99A/M102Q Set benchmark data

| ligand | PC CID<sup>k</sup> | 2D | SMILES | $\Delta G^a$ (kcal/mol) | PDB code | reference |
| --- | --- | --- | --- | --- | --- | --- |
| toluene<sup>b</sup> | 1140 |  | Cc1ccccc1 | -4.93 | ND<sup>c</sup> | [188] |
| phenol | 996 |  | c1ccc(cc1)O | -5.24 | 1LI2 [188] | [188] |
| catechol<sup>h</sup> | 289 | | c1ccc(c(c1)O)O | -4.16 ± 0.03 | 1XEP [69] | [188] |
| 2-ethoxyphenol<sup>d</sup> | 66755 | | CCOc1ccccc1O | -4.02 ± 0.03 | 3HU8 [14] | [14] |
| benzyl acetate<sup>e,f</sup> | 8785 | | CC(=O)OCc1ccccc1 | -4.48 ± 0.16 | 3HUK [14] | [14] |
| 4,5,6,7-tetrahydroindole<sup>f</sup> | 57452536 | | c1c[nH]c2c1CCCC2 | -4.61 ± 0.09 | 3HUA [14] | [14] |
| n-phenylglycinonitrile<sup>g</sup> | 76372 | | c1ccccc1NCC#N | -5.52 ± 0.18 | 2RBN [14] | [14] |
| 3-chlorophenol | 7933 |  | c1cc(cc(c1)Cl)O | -5.51 | 1LI3 [188] | [188] |
| 2-methoxyphenol | 460 |  | COc1ccccc1O | NB<sup>i</sup> | ND<sup>c</sup> | [14] |
| 4-vinylpyridine | 7502 |  | C=Cc1ccncc1 | NB<sup>i</sup> | ND<sup>c</sup> | [188] |

<sup>a</sup>T=283K, with measurements done at pH 6.8 in 50 mM potassium phosphate, 200 mM potassium chloride buffer in the case of [14]; <sup>b</sup> included for symmetry with the L99A site since this (unlike phenol and benzene) binds in both; <sup>c</sup> not determined; <sup>d</sup> fails to make crystallographic hydrogen bond [14]; <sup>e</sup> multiple binding modes; <sup>f</sup> induces helix F motion; <sup>g</sup> induces flip of Val111 sidechain; <sup>h</sup> induces flip of Leu118 sidechain; <sup>i</sup> nonbinder; <sup>k</sup> PubChem compound ID.

#### 2. Computational challenges posed by the T4 lysozyme benchmarks

Early work on the lysozyme sites focused on the difficulty of predicting binding modes [14, 118, 120] because of the slow interconversions noted above. Docking methods often can generate reasonable poses spanning most of the important possibilities [14, 70, 118, 120] but do not accurately predict the binding modes of individual compounds [14, 70, 120]. Thus, binding calculations must explore multiple potential binding modes, especially as some ligands actually populate multiple poses at equilibrium [14]. In a number of studies, candidate binding modes from docking are relaxed with MD simulations, then clustered to select binding modes for further study. It turns out an effective binding free energy for each distinct candidate binding mode can be computed separately [118] and combined to find the population of each binding mode and determine the overall binding free energy. However, this is costly, since each binding mode requires a full binding free energy calculation.

Relative binding free energy calculations do not dramatically simplify the situation,as introduction of a ligand modification can leave the binding mode uncertain. For example, adding a chlorine to phenol generates at least two binding mode variants, even if the binding mode of phenol is known) [14]. A naïve solution is to consider multiple possible binding modes in relative free energy calculations [14], but this generates multiple results; determining the true relative binding free energy requires additional information [121]. Enhanced sampling approaches provide one possible solution to the binding mode problem. For example, λ or Hamiltonian exchange techniques can incorporate sampling on an artificial energy surface where the ligand does not interact with the protein, and thus can readily reorient. Thus, approaches employing this strategy can naturally sample multiple binding modes [57, 184].

While sidechain sampling has been a significant challenge, it is possible to use biased sampling techniques such as umbrella sampling to deliberately compute and include free energies of sampling slow sidechain rearrangements [119]. However, this is not a general solution, since it requires knowing what sidechains might rearrange on binding and then expending substantial computational power to map the free energy landscapes for these rearrangements. An apparently better general strategy is including sidechains in enhanced sampling regions selected for Hamiltonian exchange [82, 91] or REST2 [186], **allowing sidechains to be alchemically softened or torsion barriers lowered (or both), to enhance sampling at alchemical intermediate states**. Reduction of energy barriers at intermediate λ states in this scheme allows enhanced conformational sampling at these λ values. Then, with swaps between λ values, enhanced sidechain sampling at intermediate states can propagate to all states, improving convergence [82, 186].

Larger protein conformational changes in lysozyme have received less attention, partly because until very recently they seemed to be a peculiar oddity only rarely observed; i.e., for ligands 4,5,6,7-tetrahydroindole and benzyl acetate in the polar site [14]. However, recent work noted above highlighted how a helix in the apolar cavity can open to accommodate larger ligands [111]. Timescales for this motion appear to be on the order of 50 ns, so it can pose sampling challenges, even for relative free energy calculations [101]. Including part of the protein in the enhanced sampling region via REST2 (described above) provides some benefits [101], but sampling these motions will likely prove a valuable test for enhanced sampling methods.

Accounting for water exchange into and out of these buried binding sites could also be a challenge in the polar binding site, as phenol binds with one ordered water [188]. However, to our knowledge, the simulation timescales for water sampling have not yet been examined, other than noting that water does not enter the polar binding site as ligands are removed [14]. In contrast, the apolar cavity is dry, both with and without a bound ligand, so water sampling is unlikely to be a challenge in this case.

## IV. FUTURE BENCHMARK SYSTEMS

Although the benchmark systems detailed above are useful, more systems are needed to expand the dataset, broaden the range of challenges, and bridge to biomedically relevant protein-ligand binding. The following subsections discuss additional systems that have already been used to test computational methods and that may be suited for development as new benchmark systems in the near future.

### A. Host-Guest Systems

The cyclodextrins (CDs; Section III A) are a particularly promising source of additional host-guest benchmark sets. Cyclic glucose polymers, the CDs are produced from starch by an enzymatic process, and are available in gram quantities at low cost from multiple suppliers. Many experimental binding data, for varied guest molecules, are available for this class of host molecules, especially for 6-membered α-CD and 7-membered β-CD, which both have adequate aqueous solubility. Indeed, a thorough review 1998 review tabulates hundreds of binding data [145], and many additional measurements have been published since then; e.g. references [21, 32, 33, 112, 149, 162, 193]. A number of these studies were done calorimetrically, and thus provide not only binding free energies but also binding enthalpies. Fewer data are available for 8-membered γ-CD, and the greater diameter of this host makes for a less well-defined and floppier binding cavity. Overall, the amount of existing binding data for CDs greatly exceeds what is currently available for CB7 and other cucurbiturils.

Structurally, the binding cavity of β-CD is about the same size as CB7 (Section IIIA1), but the cyclodextrins are more flexible than CB7, as their glucose monomers are joined by one single bond, whereas CB7's glycouril monomers are joined by two single bonds. The CDs also appear easier to derivatize than the cucurbiturils. In particular, varied substituents may be appended by reactions involving secondary hydroxyls at the wide entry and primary hydroxyls at the narrow entry [41, 53, 85, 142, 171], with the caveat that generating pure products can be difficult, because there are so many hydroxyls that may be modified. Binding data on CD derivatives could be quite useful as a means of adding chemical diversity to host-guest benchmark sets, and such data are already available for some derivatives (see, e.g., [46, 145]).

The CDs are computationally tractable [9, 16, 23, 61, 73, 92, 107, 108, 154, 156, 190, 191, 202], and thus can be used for “hard” benchmark sets to test force fields (Section II A 1 c). One complicating feature for calculations, relative to CB7, is that the two entryways to a CD are not equivalent, and asymmetric guest molecules may prefer to bind “head-in” or “head-out”. As these two binding modes typically do not interchange on the microsecond timescale, they must be considered separately when one computes binding thermodynamics [73]. It is also worth noting that, because the CDs are glucose polymers, they may be best modeled with dedicated carbohydrate force fields [22, 72, 94, 195], rather than generalized small molecule force fields. A number of CD (and other host guest) binding systems are available for download in electronic format at the BindingDB [67, 106] website (www.bindingdb.org/bind/HostGuest.jsp)

### B. Protein-Ligand Systems

On the order of a million experimental proteinligand binding measurements are currently accessible through open-access databases, notably BindingDB [106], ChEMBL [10] and PubChem [93]. These databases can be valuable sources of, or at least starting points for, new protein-ligand benchmark datasets. In fact, automated procedures have already been used to extract about 700 downloadable validation sets (www.bindingdb.org/validation_sets/index.jsp), each comprising a congeneric series of 10-50 ligands with binding data against a defined protein, and a cocrystal structure for at least one of the ligands in the series. However, there is still a need for a smaller collection of highly optimized benchmark sets as research foci for the computational chemistry community. Such sets should exemplify specific challenges not well covered by existing benchmark systems; be based upon high quality binding measurements for at least 20 ligands, measured by consistent procedures across all ligands; and include crystal structures for the apo-protein and cocrystal structures for multiple ligands. Analysis and extraction of such sets from the big databases is a promising future direction. Here, however, we take the less systematic but more expedient approach of considering several systems that have already proven themselves to be computationally tractable and informative.

#### 1. Constructed Binding Sites in Cytochrome C Peroxidase

Two artificial binding sites have been designed into the enzyme cytochrome C peroxidase (CCP): the “closed” site, created by the mutation W191G [49, 147]; and the “open” site mutant, created by supplementing mutation W191G with partial deletion of a loop, and thus opening the site to the outside of the protein [134, 147, 151]. As for the artificial binding cavities in T4 lysozyme (Section III B), the two CCP sites bind simple, fragment-like ligands with modest binding free energies (e.g., −3 to −7 kcal/mol for the open site [147]); discovery of new ligands is relatively straightforward [15, 147]; and new crystal structures can be obtained fairly readily. The protein has relatively modest size (around 280 residues) as in, for example, the 1KXM structure.

What is new is that the CCP binding sites contain an ASP which appears to be charged, at least in the presence of some ligands, as evidenced by observed interactions in structures, and by experiments where replacing the ASP with ASN abolishes binding of imidazoles, despite minimal changes to the binding site structure [50]. This side-chain interacts at short range with the ligands, which may be cationic or polar neutral compounds. As a consequence, the CCP sites challenge the ability of computational methods to accurately account for strong electrostatic interactions in the low-dielectric interior of a protein [147]. Indeed, free energy calculations with two different force fields and distinct computational methods were found to overestimate the range of affinities, across a series of ligands, about three-fold, for the closed site [6, 7], and a similar pattern appears to hold for the open site [147]. Further analysis suggests that these overestimates stem from overestimated electrostatic interactions in these buried sites [147], perhaps because the force fields used did not account explicitly for electronic polarization. Additionally, the fact that different CCP ligands have different net charges may pose methodological challenges for some free energy techniques [148]. One modeling challenge is that CCP contains a heme, providing additional setup challenges (though since this is such an important cofactor, these may be worthwhile to face). In summary, these model CCP binding sites appear challenging yet tractable, and have already yielded insight regarding possible directions for force field improvements. Thus, they are good candidates to provide new benchmark sets in the near future.

#### 2. Thrombin

Human thrombin, an enzyme of about 300 residues, is interesting both as a drug target related to blood coagulation and as a representative serine protease. Binding and structural data are available for a wide variety of ligands ([8, 141, 168, 177] and others), and have already been used as the basis for free energy simulations [17, 186, 187]. Some of the experimental binding data, obtained calorimetrically, highlight interesting trends, such as non-additivity (positive coupling) between substitutions at different locations on the chemical scaffolds of two compound series [8].

One computational study obtained relative free energies for two compound series that largely captured experimental trends and achieved an overall mean unsigned error of 0.74 kcal/mol over a range of roughly 5 kcal/mol. However, the accuracy of the results differed significantly between the two series [17]. Another relative binding free energy study for thrombin found that the results changed, depending on the initial conformation of the ligands, and showed that enhanced sampling techniques could reduce the dependnce on starting conformation [186]. These prior studies suggests thrombin may be at a “sweet spot” for benchmark systems, where the system is relatively tractable, and encouraging results have been obtained, but where there are still clear challenges.

#### 3. Bromodomain proteins

Bromodomains (BRDs) are a family of protein domains of about 100 residues that bind actylated lysine residues at the surface of histones and thus read out epigenetic markers. Bromodomains are present in many human proteins, and are being explored as potential drug targets for diseases including cancer and atherosclerosis [2]. These compact domains can be expressed, purified and studied as independent proteins, and are associated with a growing body of small molecule binding data [5, 28, 48, 68, 75, 143, 196, 201], including some obtained by isothermal titration calorimetry [28, 48, 68, 75].

The small size of bromodomains, and the fact that their binding sites are relatively solvent-exposed, makes them particularly suitable for free energy simulations. One recent study, which used alchemical techniques to compute absolute binding free energies of 11 different ligands for bromodomain BRD4 [2], achieved a remarkable level of accuracy, RMS error 0.8 ± 0.2 kcal/mol, for binding free energies spanning a range of 4 kcal/mol. Docking calculations included in the same study did not work as well. The compounds studied were diverse, and therefore not amenable to relative free energy methods in which one ligand is computationally converted into another. Overall, then, this appears to be class of systems that could yield relatively tractable and informative proteinligand benchmark systems. One known challenge is that some ligands have multiple plausible binding modes [2]. In addition, a diverse ligand series can pose severe challenges for relative free energy techniques.

#### 4. Other protein-ligand systems

Several other systems may be of possible interest because of the wealth and quality of experimental data, the extent of prior computational work, or the combination of pharmaceutical relevance with important challenges. None of these seem to be well-characterized or well-studied enough yet to be benchmark systems, but they may be interesting choices for the future. The first system we would put in this category is trypsin, and especially binding of benzamidine and its derivatives. While trypsin has been the focus of several free energy studies [36, 83, 84, 173, 181] and the SAMPL3 challenge [135, 164], it also appears to be subject to extremely slow protein motions (on the tens of microsecond timescale) which are only beginning to be characterized [140]. As a consequence, short free energy calculations may appear to converge [83, 84, 173], but longer calculations can reveal a profound lack of convergence [140]. Therefore, trypsin may become a good benchmark for studying binding that is coupled with slow conformational dynamics; enhanced sampling methods and Markov state models [140] may be particularly helpful here.

HIV integrase is also of potential interest as a benchmark system. Indeed, blind predictions for a set of fragments binding to several sites formed part of the basis of the SAMPL4 challenge [122, 139]. Although the range of affinities in this set is modest (200 to 1450 μ*M* [139]), many crystal structures are available, as is a wealth of verified nonbinders [139]. One group actually did remarkably well using free energy calculations to recognize binding modes and predict nonbinders [56, 122]. Coupled with other HIV integrase data available in the literature and binding databases for larger ligands or other series, this may make this system attractive for future benchmarks.

The protein FKBP has also been the focus of several different free energy studies on the same series of ligands over the years [54, 55, 81, 99, 157, 183, 200]. However, in many cases, differences in calculated values between different studies-even with what appears to be the same force field and system preparation–are larger than differences relative to experiment [54, 81]. Some challenges are clear, such as conformational sampling for some of the larger ligands [157], and there have been some suggestions of force field issues and long equilibration times for the protein [55]. Perhaps this may be suitable as a benchmark system in the near-term as well.

A number of other proteins also have strong potential to generate useful benchmark sets. For example, free energy calculations have been carried out for influenza neuraminidase inhibitors [166] with some success [115], but other series include more complicated structure-activity relationships and are associated with protein loop motions that may be difficult to model [90]. Periplasmic oligopeptide binding protein A (OppA) binds a series of two to five-residue peptides, for which there exists a large amount of calorimetric and crystallographic binding data [35, 165, 174], and the system appears challenging but potentially tractable for free energy calculations [110]. The JNK kinase may pose an interesting conformational sampling challenge, due to its slow interconversion between binding modes for some ligands [89]. And the ongoing series of Drug Design Data Resource (D3R) [] blinded challenges may also be source of informative protein-ligand systems (drugdesign-data.org/about/datasets) that are familiar to the computational chemistry community. As noted above, however, many other protein-ligand binding systems have been characterized experimentally, and a systematic filtering would undoubtedly yield more benchmark candidates.

## V. HOW TO USE BENCHMARK SYSTEMS

Benchmark systems will have multiple uses, as discussed above, but not all benchmark systems can cover all uses. Some will be particularly valuable for testing accuracy relative to experiment. For this purpose, relatively large numbers of ligands are needed to afford meaningful statistics. Other benchmark systems will be more useful for testing sampling techniques, and still others will, at least initially, serve as test beds to determine the sensitivity of computational results to various factors.

In our view, benchmark systems will serve also to help design careful computational experiments. For example, researchers can test whether a particular method is sampling the motions which others have already shown to be important, or how the choice of starting conformation impacts the rate of convergence to a known, gold standard value for a particular force field and system composition. The availability of benchmark systems will also facilitate comparisons where only a single piece of a workflow is modified. For example, one may ask how results change if a different protonation state assignment tool is used to prepare a protein. Of course, such comparisons can already be done, but the results will be far more useful in the context of generally accepted and widely used test cases.

We hope that, ultimately, results from reliable “gold standard” binding free energy computations will be available for a set of benchmark systems. These would be from fully converged binding free energy calculations, and give correct results for a particular force field and system preparation, allowing quantitative comparison of the force field results with experiment. Such results will also facilitate a great deal of science on method efficiency, as new methods which purport to be more efficient could easily and *automatically* be run on a standard set of systems to see how much more efficient they are than the (perhaps brute-force) method which yielded the gold standard results. Thus, various enhanced sampling methods could easily be observed to have strengths and weaknesses on known problem classes. These systems will allow automated testing of the efficiency of new methods on real-world problems.

## VI. WE NEED WORKFLOW SCIENCE

While the benchmark systems discussed here will already be useful, to fully realize their benefits a great deal of engineering needs to be done to facilitate workflow science. Currently, a wide variety of computational tools are available for different stages of the free energy calculation process, from system preparation (protonation state assignment, building in missing residues and loops, adding counterions, etc.) to force field assignment, to planning and conducting the calculations themselves (choice of method, simulation package, and so on). Often, each set of tools lives in its own ecosystem and is not designed to be easily interchangeable with tools from another ecosystem. This makes it very difficult to systematically compare methods that differ only in one respect; instead, one must adopt an entirely different toolset to change one aspect of a procedure. For example, swapping different tools for assigning protein protonation states could yield valuable insights into the relative merits of these tools and the importance of protonation at specific residues, but currently, this is, at best, an arduous task.

### A. Workflow automation is needed

At the most basic level, we need to allow calculations to be easily repeated on all of the benchmark systems via automated workflows. One should not have to become an expert in the systems being studied in order to be able to successfully apply calculations to them; inputs should be easily available and repeating calculations should become fully automated so that a new method can be tested by simply specifying the set of benchmarks to run on.

To achieve this, at least two major innovations are needed. First, we need automated workflows that can proceed from the specification of a system to target to yielding the desired results without human intervention. Second, we need a standard data structure for input to and output from these workflows so that people can easily obtain inputs for benchmark systems and only change the component they want to change (such as the force field or system preparation) and leave the other components unchanged so that, in an automated manner, they can focus their testing on only the components they want to test.

### B. Analysis automation will also be needed

At the most basic level, we can simply check whether we are getting the expected answer for each calculation performed, at least for systems where a gold standard result is available. However, this does not provide nearly enough insight, especially in cases of failure, where we would like insight into *why* we failed. Are the relevant motions being sampled? Do we have the right protonation states and binding modes? Many other factors may need to be considered. We need ways to automatically check that we are sampling the right motions, identifying correct binding modes/conformations, and so on, without having to become experts on the specific systems examined. Probably we will need to define ways to automatically specify what order parameters should be monitored to assess for adequate sampling.

### C. Modularization will be key

To achieve these goals, researchers should develop or package tools so that they take a set of well specified inputs and provide well specified outputs in an interchangeable way. This may involve containerizing key pieces of workflows such as in Docker [40] or Singularity [95] containers, and developing standards as to what inputs and outputs are provided to each component of the workflow. Another key goal of modularization is to separate the *operator* from the *method*. Currently, binding calculations are most often done by a human expert, who makes a variety of decisions along the way (though Schrodinger's workflow represents real progress toward automation [187]), making it difficult to separate the importance of human expertise from the merits of the methods employed. Containerizing and modularization will be key for this, allowing methods to be employed only in a well-defined way which is reproducible. It is this type of science–coupled with benchmark tests–which is needed to advance the field.

## VII. THE FUTURE OF THIS WORK

This work has so far presented a small group of benchmarks for binding free energy calculations, highlighted some of the ways in which they have already proven their utility, and has suggested some potential future benchmark systems. We hope that the community will become involved in identifying, characterizing, and helping to select additional benchmark systems, both from those proposed here as well as from systems which are currently being studied or which we have overlooked. We seek community input to help characterize, identify and share such systems. We also expect that there may be community interest in test systems specifically selected to challenge sampling algorithms, without reference to experimental data.

In order to provide for updates of this material as new benchmark systems are defined, and to enable community input into the process of choosing them, we have made the LaTeX source for this article on GitHub at http://www.github.com/mobleylab/benchmarksets, with each version having a permanent DOI assigned by Zenodo. We encourage use of the GitHub issue tracker for discussion, comments, and proposed updates. We plan to incorporate new material via GitHub as one would for a coding project, then make it available via a preprint server, likely bioRxiv. Given substantial changes to this initial version of the paper, it may ultimately be appropriate to make it available as a “perpetual review” [123] via another forum allowing versioned updates of publications.

Ideally, we might also update this work in the future with results from “standard” calculations on the benchmark systems discussed here, along with links to code to allow reproduction of those calculations.

## VIII. CONCLUSIONS AND OUTLOOK

Binding free energy calculations are a promising tool for predicting and understanding molecular interactions and appear to have enough accuracy to provide substantial benefits in a pharmaceutical drug discovery context. However, progress is needed to improve these tools so that they can achieve their potential. To achieve steady progress, and to avoid potentially damaging cycles of enthusiasm and disillusionment, we need to understand and be open and honest about key challenges. Community adoption of well-chosen benchmark systems is vital for this, as it will allow researchers to rigorously test and compare methods, arrive at a shared understanding of problems, and measure progress on well-characterized yet challenging systems. It is also worth emphasizing the importance of sharing information about apparently well thought-out and even promising methods that do *not* work, rather than sharing only what does appear to work. Identifying and addressing failure cases and problems is critically important to advancing this technology, but failures can be harder to publish, and may even go unpublished, even though they serve a unique role in advancing the field. We therefore strongly encourage that such results be shared and welcomed by the research community. Potentially, the GitHub repository connected with this perpetual review paper could serve as a place to deposit input files and summary results of tests on these benchmark systems, with summary information perhaps being included in this work itself or topical sub-reviews within the same repository.

Here, we have proposed several benchmark systems for binding free energy calculations. These embody a subset of the key challenges facing the field, and we plan to expand the set as consensus emerges. Hopefully, these systems will serve as challenging standard test cases for new methods, force fields, protocols, and workflows. Our desire is that these benchmarks will advance the science and technology of modeling and predicting molecular interactions, and that other researchers in the field will contribute to identifying new benchmark sets and updating the information provided about these informative systems.

## Acknowledgments

DLM appreciates financial support from the National Institutes of Health (NIH; 1R01GM108889-01) and the National Science Foundation (NSF; CHE 1352608). MKG thanks the NIH for partial support of this work through grants R01GM061300, R01GM070064 and U01GM111528. The contents of this publication are solely the responsibility of the authors and do not necessarily represent the official views of the NIH or the NSF.

We also appreciate helpful discussions with a huge number of people in the field, including a wide variety of participants at recent meetings such as the 2016 Workshop on Free Energy Methods in Drug Discovery. Conversations with John Chodera (MSKCC), Chris Oostenbrink (BOKU), Julien Michel (Edinburgh), Robert Abel (Schrödinger), Bruce Gibb (Tulane), Matt Sullivan (Tulane), and Lyle Isaacs (Maryland) were particularly helpful.

